# Social behavior and anxiety contribute to nicotine self-administration in adolescent outbred rats

**DOI:** 10.1101/257097

**Authors:** Tengfei Wang, Wenyan Han, Apurva Chitre, Oksana Polesskaya, Leah C. Solberg Woods, Abraham A. Palmer, Hao Chen

## Abstract

Both emotional and social traits interact with genetic factors to influence smoking behavior. We previously established a socially acquired nicotine intravenous self-administration model where social learning of a nicotine-associated odor cue reversed conditioned flavor aversion and promoted nicotine intake. In this study, we first phenotyped ~ 800 adolescent heterogeneous stock rats in open field, novel object interaction, social interaction, elevated plus maze, and marble bury behaviors. These rats were then phenotyped on socially acquired nicotine self-administration. We found 243 significant correlations between different behavioral tests. Principal component regression analysis found that ~ 10–20% of the variance in nicotine-related measures, such as intake during the first or the last three fixed-ratio sessions, the progressive ratio session, and reinstatement behavior, can be explained by variations in behavioral traits. Factors corresponding to social behavior and anxiety were among the strongest predictors of nicotine intake and reinstatement of nicotine-seeking behavior. We also found many sex differences in behavioral measures. These data indicated that the genetic diversity of this population, in combination with social behavior and anxiety, are significant contributors to the divergent nicotine self-administration behavior and indicated a high probability of discovering sex-specific genetic mechanisms for nicotine intake in future genome-wide association studies.

## Introduction

Smoking behavior is affected by a myriad of environmental and genetic factors. The strong influence of social environment in smoking initiation, especially during adolescence, the most critical age for smoking initiation^1^, is well-documented^2,3^. Many personality and affect-related traits are also known to influence smoking behavior^4^, such as anxiety^5,6^, negative affect^7^, or response to stress^8^. Genetic factors are also critical to cigarette smoking^9,10^. The overall heritability of smoking is estimated to be 0.4 – 0.6 in human^11,12^. In addition, genetic variants of many genes, such as the CHRNA5-CHRNA3-CHRNB4 cluster^13^^-^^15^, CYP2A6^16^, contribute to smoking behaviors. However, less than 1% of the heritable variance in smoking behaviors was attributed to these polymorphisms, indicating that most of the genetic factors determining smoking behaviors are still unknown^13,15^.

We have developed a rat model of adolescent nicotine self-administration that captures the role of social learning in promoting nicotine intake. This operant licking model delivers intravenous nicotine with a contingent olfactogustatory cue. Our basic finding was that rats trained alone under this condition developed conditioned flavor aversion and failed to self-administer nicotine. However, interacting with another rat that consumed the same olfactogustatory cue during self-administration reversed the conditioned aversive response and promoted nicotine intake^17^. We have further observed that the presence of a nicotine-associated olfactory cue^17^ and carbon disulfide^18^, a component of rodent breath, are two critical signals for the effect of social learning on nicotine intake. In contrast, the gustatory cue did not appear to be involved^17^. In fact, IVSA (as it hasn’t been defined in the manuscript yet) could be established even when an aversive gustatory cue was used^19^.

This socially acquired nicotine self-administration model provided us with an opportunity to delineate the contributions of various environmental, emotional, and genetic factors to adolescent-onset nicotine self-administration. By assessing behaviors related to locomotion, novelty object interaction, anxiety-like phenotype, and social interaction in each rat before nicotine self-administration, we can correlate aspects of these behavioral traits with nicotine intake. Using inbred strains of rats, we have estimated the heritability of nicotine intake to be 0.54 – 0.65 in this model^20^. Thus, the contribution of genetic factors to nicotine intake can also be studied.

Here we summarize data obtained from approximately 800 adolescent heterogeneous stock (HS) rats. The HS rats were derived from eight inbred founder strains^21^. After more than 70 generations of rotational out-breeding, each HS rat represents a unique random mosaic of the founder haplotypes^22^. This population has been successfully used to map many physiological and behavioral traits^23^^-^^27^ Our results showed that features extracted from the non-nicotine behavioral profiling explained 10–20% of the variation in nicotine intake.

## Results

### Breeding and study population

A total of 8 batches of rats were processed between 2014-2017. Each batch started with 25 male and 25 female non-sibling breeders. Breeding pairs were assigned to maximize genetic diversity of the population. The average litter size was 9.1±0.21. Litters were culled to a maximum of 8 pups to ensure standard nutrition. Rats were weaned on postnatal day (PND) 21. A radio frequency identification chip (RFID) was inserted subcutaneously when rats were weaned. Two male and two female rats per litter were used for behavioral studies. No significant variations in litter size were observed between batches (F_7, 158_ = 2.0, *p* > 0.05).

### Open field test (OFT)

From each litter, two males and two females rats were randomly selected to receive five behavioral tests starting on PND 32, one test per day, starting from the OFT (see Table 1). Rats were placed in a 1 m × 1 m square arena for 1 h during this test. As shown in Table 2, female rats traveled a greater distance (1.07 ×) than did the males in the arena. Females also stayed slightly closer (0.98 ×) to the center of the arena, entered the center zone more frequently (1.22 ×), and spent more time in the center zone than did the males (1.27 ×). However, there was no difference in the latency of entering the center zone between the female and male rats. Taken together, female adolescent HS rats explored the open field, including the center zone, more than the males.

**Table 1.** Schedule for behavioral tests

| PND | Procedure |
| --- | --- |
| 32 | Open field |
| 33 | Novel object |
| 34 | Social interaction |
| 35 | Elevated plus maze |
| 36 | Marble bury |
| 38 | Surgery & recovery |
| 41 – 50 | Nicotine IVSA, FR |
| 51 | Nicotine IVSA, PR |
| 52 – 60 | Extinction, Reinstatement |

**Table 2.** Open field test summary statistics

|  | n | Travel Distance<br>Total (cm) | Distance to Center<br>Mean (cm) | Center<br>Duration (s) | Center<br>Frequency | Center<br>Latency (s) |
| --- | --- | --- | --- | --- | --- | --- |
| F | 387 | 14636.4 ± 3846.3 | 33.3 ± 3.5 | 65.9 ± 59.0 | 21.0 ± 14.9 | 277.9 ± 824.8 |
| M | 377 | 13688.1 ± 3774.7 | 34.1 ± 3.4 | 52.0 ± 50.2 | 17.2 ± 13.9 | 274.9 ± 375.6 |
| F/M ratio |  | 1.07 | 0.98 | 1.27 | 1.22 | 1.01 |
| Effect size |  | 0.25 | 0.22 | 0.24 | 0.25 | 0 |
| p |  | 0.0006 | 0.003 | 0.0009 | 0.0007 | 0.99 |

### Novel object interaction test (NOIT)

The NOIT was conducted in the same arena as the OFT with the exception that a cylindrical cage was placed in the center of the arena. Compared to the OFT, rats stayed significantly closer to the center of the arena (F_1,1406_ = 1155.6, p < 2*e* – 16), visited arena center more frequently (F_1,1406_ = 208.0, *p* < 2*e* – 16), spent significantly more time in the center of the arena (F_1, 1406_ = 50.7, *p* = 1.7*e* – 12), and entered arena center with shorter latency (F_1,1406_ = 866.7, *p* < 2*e* – 16) during the NOIT. These data indicated that placing an novel object in the center of the arena changed the behavior of the rats, and thus this test measured a different behavioral process compared with the OFT.

Consistent with the OFT, female rats traveled a greater distance than male rats (1.07 ×). The females also stayed closer to the center zone (0.92 ×), spent more time in the center zone (1.13 ×), and entered the center zone sooner (0.79 ×) compared with male rats. However, there was no sex difference in the frequency of entering the center zone (Table 3). These results demonstrated that adolescent female HS rats had more interaction with the novel object than did the males.

**Table 3.** Novel object test summary statistics

|  | n | Travel Distance<br>Total (cm) | Distance to Center<br>Mean (cm) | Center<br>Duration (s) | Center<br>Frequency | Center<br>Latency (s) |
| --- | --- | --- | --- | --- | --- | --- |
| F | 381 | 7684.3 ± 1998.8 | 23.5 ± 5.7 | 276.7 ± 172.1 | 33.8 ± 18.9 | 83.6 ± 131.9 |
| M | 371 | 7204.3 ± 1990.7 | 25.5 ± 6.7 | 244.8 ± 174.7 | 32.4 ± 22.2 | 105.6 ± 149.6 |
| F/M ratio |  | 1.07 | 0.92 | 1.13 | 1.04 | 0.79 |
| Effect size |  | 0.23 | 0.32 | 0.18 | 0.07 | 0.16 |
| p |  | 0.002 | <1e-04 | 0.01 | 0.37 | 0.03 |

### Social interaction test (SIT)

Two identical cylindrical rat enclosures were placed in the arena, one of them housed a weight matched Sprague-Dawley rat of the same sex as the HS subject rat, while the other was empty. The HS rats paid more visits (72.55 ± 32.52 vs 37.55±17.39, *p* < 2.2*e* – 16), waited less time before entering (4.05±21.13 vs 38.04±56.26s, *p* < 2.2*e* – 16), stayed longer (440.25±204.64 vs 138.59±84.8s, *p* < 2.2*e* – 16), and remained closer (11.74±4.64 vs 23.91±5.46cm, *p* < 2.2*e* – 16) to the social zone than the object zone. The was no sex difference in total travel distance, distance to the object zone, latency of entering either the object or the social zone, or frequency of entering the social zone. However, female rats remained closer to the social zone (0.90 ×), spent less time in the object zone (0.80 ×) but more time in the social zone (1.18 ×), and paid fewer visits to object zone (0.81 ×) compared with male rats (Table 4). These data indicated that adolescent HS rats were pro-social and that the females had stronger tendency to engage in social interaction with unfamiliar rats than did males.

**Table 4.** Social interaction test summary statistics

|  | n | Travel Dist<br>Total (cm) | Dist to Obj Zone<br>Mean (cm) | Dist to Soc Zone<br>Mean (cm) | Obj Zone<br>Duration (s) | Obj Zone<br>Frequency | Obj Zone<br>Latency (s) | Soc Zone<br>Duration (s) | Soc Zone<br>Frequency | Soc Zone<br>Latency (s) |
| --- | --- | --- | --- | --- | --- | --- | --- | --- | --- | --- |
| F | 381 | 8716.3 ± 2262.9 | 23.7 ± 5.3 | 11.1 ± 5.0 | 123.1 ± 81.1 | 33.7 ± 15.7 | 38.8 ± 58.9 | 475.7 ± 211.0 | 71.8 ± 34.5 | 3.9 ± 14.9 |
| M | 370 | 8985.1 ± 2248.4 | 24.2 ± 5.6 | 12.4 ± 4.1 | 154.5 ± 85.7 | 41.5 ± 18.2 | 37.3 ± 53.4 | 403.7 ± 191.4 | 73.3 ± 30.4 | 4.2 ± 26.1 |
| F/M ratio |  | 0.97 | 0.98 | 0.9 | 0.8 | 0.81 | 1.04 | 1.18 | 0.98 | 0.93 |
| Effect size |  | 0.12 | 0.09 | 0.29 | 0.38 | 0.46 | 0.03 | 0.36 | 0.05 | 0.02 |
| p |  | 0.10 | 0.21 | <1e-04 | <1e-04 | <1e-04 | 0.72 | <1e-04 | 0.53 | 0.82 |

### Elevated plus maze test (EPM)

HS rats were tested in a standard elevated plus maze in the next session. Overall, HS rats visited more frequently (16.25±8.10 vs 10.54 ±7.03, *p* < 2.2*e* – 16), spent more time (225.75±67.32 vs 63.67± 46.86s, *p* < 2.2*e* – 16), and waited a shorter period of time (36.77±43.91 vs 99.76±113.00s, *p* < 2.2*e* – 16) before entering the closed arm than the open arm. There was no sex difference in the frequency and latency of entering the open arm, the closed arm, or the center zone. However, female HS rats traveled a greater distance (1.05 ×), spent more time in the open arm (1.26 ×) and the center zone (1.13 ×), and thus less time in the closed arm (0.93 ×) than did the males (Table 5). These results indicated that adolescent male HS rats have higher anxiety than females.

**Table 5.**
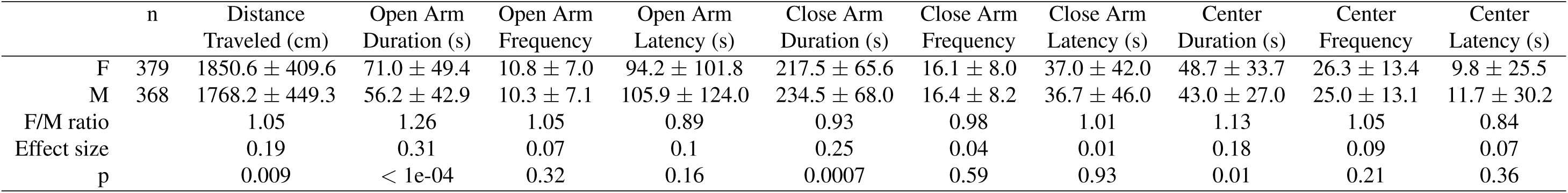
Elevated plus maze summary statistics

### Marble bury test (MBT)

The final behavioral test was the Marble bury test, where rats were placed in an arena with 20 marbles laid out as a 5 × 4 grid for 6 min. Rats started to dig the bedding after 84.1±93.9s, and left 7.8±4.5 marbles unburied. There was no significant sex difference in the marble bury test (Table 6).

**Table 6.** Marble bury test summary statistics

|  | n | Number Left | Dig Latency (s) |
| --- | --- | --- | --- |
| F | 382 | 8.0 ± 4.7 | 78.1 ± 96.9 |
| M | 375 | 7.7 ± 4.3 | 90.2 ± 90.5 |
| F/M ratio |  | 1.04 | 0.87 |
| Effect size |  | 0.06 | 0.13 |
| p |  | 0.40 | 0.08 |

### Correlations between behavioral tests

Pearson correlations between behavioral test scores were calculated using the corr. test function in the Psych package of R. Multiple comparisons were adjusted using the Holm method. There were 354 correlations with adjusted *p* < 0.05. Among them, 111 were from measures of the same test and the 243 remaining were from measures of different tests. These results are summarized as a heatmap (Figure 1). In general, correlations within the same behavioral test have relatively high coefficients. Details of these findings are provided in Tables S1 – S5. In comparison, most of the cross-test correlations had moderate to low coefficients. Significant correlations between behavioral tests are shown in Table S6 (for *r* > 0.2) and in Table S8 (for *r* < 0.2). Notably, total travel distances, which was measure in four of the tests descried above, showed some of the highest correlations. There were also strong correlations between the NOIT and the OFT, which used the same test arena. Measures of SIT and MBT had some of the weakest correlations with other tests, which was consistent with our expectation that they measured distinct aspects of rodent behavior.

**Figure 1.**
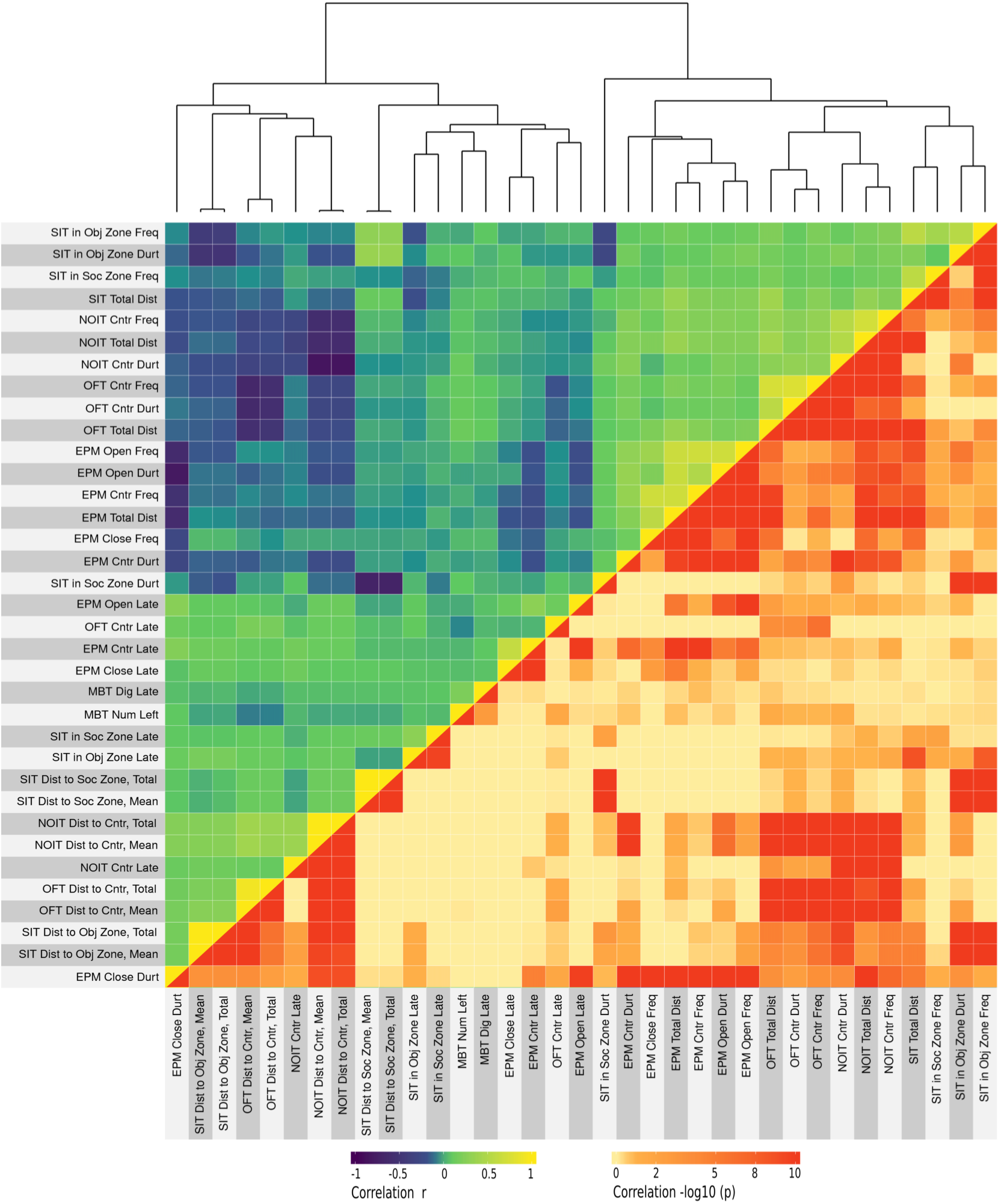
Correlations between behavioral tests. A total of 35 measures were obtained from the six behavioral test conducted before the commencement of nicotine self-administration. Pearson correlation coefficient between these measures were plotted on the upper (left) triangle of the heatmap. The Holm-adjusted p values (converted using-log10) for these correlations were shown on the lower (right) triangle of the heatmap. The top dendrogram was obtained using hierarchical clustering. OFT: open-field test, NOIT: novel object interaction test, SIT: social interaction test, EPM: elevated plus maze, MBT: marble bury test.

### Socially acquired nicotine intravenous self-administration (IVSA)

IVSA started on PND 41. The first 10 daily sessions used a fixed ratio 10 reinforcement schedule to concurrently deliver a flavor cue containing carbon disulfide to the active spout and i.v. nicotine. Carbon disulfide is a component of rodent breath and part of the complex signal that facilitated socially acquired nicotine IVSA^18^. A progressive ratio test was conducted on session 11. Catheter patency was validated thereafter. Among the 730 rats that started nicotine IVSA, 88 rats were excluded from further analysis due to experimental errors or catheter failure.

The number of licks on both spouts during the fixed-ratio sessions are shown in Figure 2. Due to the wide range of licking activity, data were split into two groups using the median nicotine infusion of the 10 fixed ratio sessions. For rats with higher nicotine infusions (Figure 2A), the number of licks on the active spout was significantly greater than those on the inactive spout (F_1,298_ = 83.9, *p* < 2*e* – 16); females licked significantly more than the males (F_1,289_ = 10.65, *p* = 0.001); the number of licks on the active spout increased significantly (F_9, 2789_ = 5.79, *p* = 5.3e-08), while the number of licks on the inactive spouts significantly decreased over the course of the sessions (F_9,2812_ = 37.9, *p* < 2*e* – 16). In contrast, for rats with lower nicotine infusions (Figure 2B), the number of licks on the active spout was significantly fewer on the active spout than on the inactive spout (F_1,304_ = 238.4, *p* < 2*e* – 16); females licked significantly more than the males (F_1,304_ = 9.63, *p* = 0.002); but the number of licks on the active (F_9, 2797_ = 35.13, *p* < 2*e* – 16) and inactive spouts (F_9, 2797_ = 43.28, *p* < 2*e* – 16) both decreased significantly over sessions. These results are consistent with our previous finding that, despite being paired with an appetitive cue, i.v. nicotine has aversive properties that are reversed by social learning, but genetic factors also plays significant roles in determining the effect of social learning^20^. The number of nicotine infusions during the last three fixed ratio sessions are shown in Figure 2C and 2D for females and males, respectively. The mean infusion distributed in a wide range [0, 30.9].

**Figure 2.**
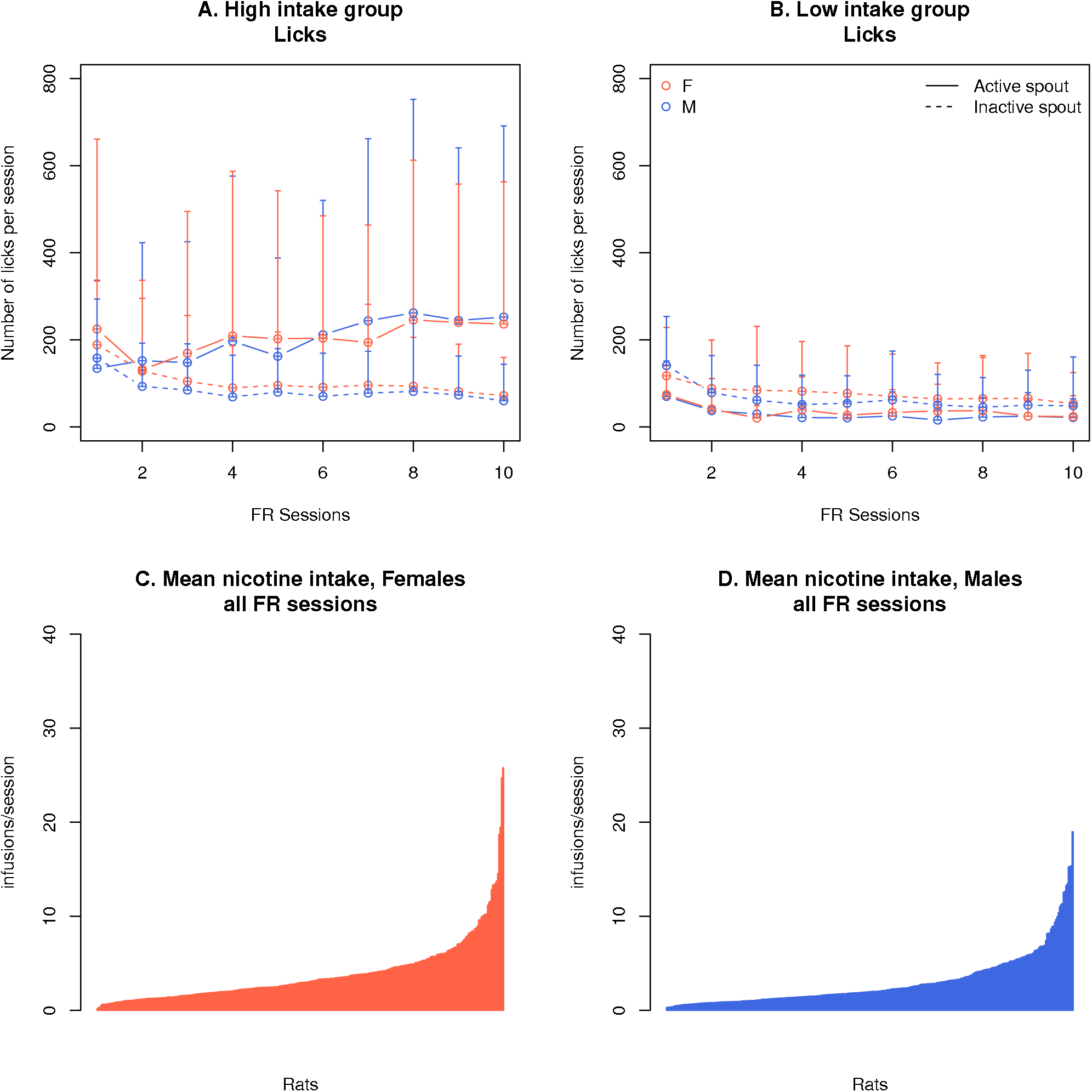
Socially acquired nicotine self-administration, fixed ratio 10. Rats were split into high vs low nicotine groups based on the median of their nicotine infusion during the 10 fixed-ratio sessions. The high intake groups licked significantly more on the active spouts compared to the inactive spout (A), while the reverse was true for the low intake group (B). Their number of total infusions were plotted in ascending order for females (C) and males (D).

Similar to other behavior tests, there were many sex differences in nicotine self-administration. For example, females received significantly more infusions than males during the first three (*p* = 0.007, Figure 3A) and the last three (*p* = 0.007, Figure 3B) fixed ratio sessions. Females also reached higher breakpoints during the progressive ratio test (*p* = 0.002, Figure 3C). Lastly, rats licked more on the active than on the inactive spout during the reinstatement test (*p* < 2*e* – 16). The number of licks on the active spout was greater in females than in males (*p* < 2*e* – 16). However, there was no sex difference in the number of licks on the inactive spouts (*p* = 0.998). Thus both males and females demonstrated contextual cue-induced reinstatement but drug-seeking was stronger in females.

**Figure 3.**
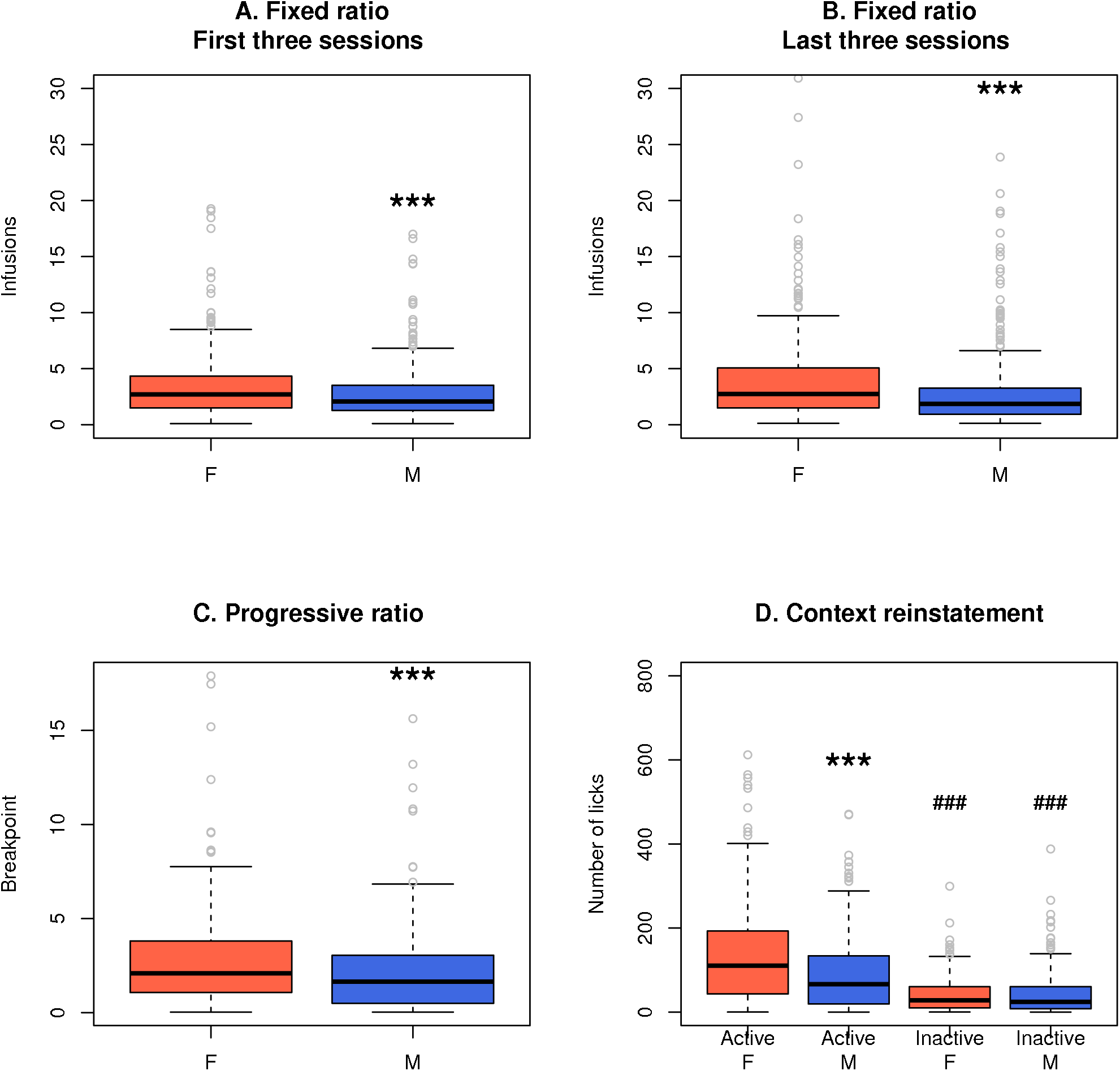
Sex differences in socially acquired nicotine self-administration. Females obtained significantly more nicotine infusions during the first three (A) and the last three (B) fixed-ratio sessions and during the progressive ratio session (C) than males. Both females and males licked significantly more on the active spout than on the inactive spout during the contextual cue-induced reinstatement session. Females also showed stronger reinstatement than males. ***: *p* < 0.001, compared to females. ###: *p* < 0.001 compared to the corresponding active spout.

### Principal component regression analysis

All data from the behavioral tests conducted before the initiation of nicotine IVSA, including OFT, NOIT, SIT, EPM, and MBT, were first scaled to [0,1] and centered. A principal component analysis was conducted, which resulted in 35 PCs. Varimax rotation was used to obtain orthogonal factors to facilitate the interpretation of these components. The amount of variance explained by these PCs were shown in Figure S1. As shown in Figure 4, PCs 1-31 all had strong loading on at least one of the behavioral measures. These PCs were retained for subsequent regression analysis.

**Figure 4.**
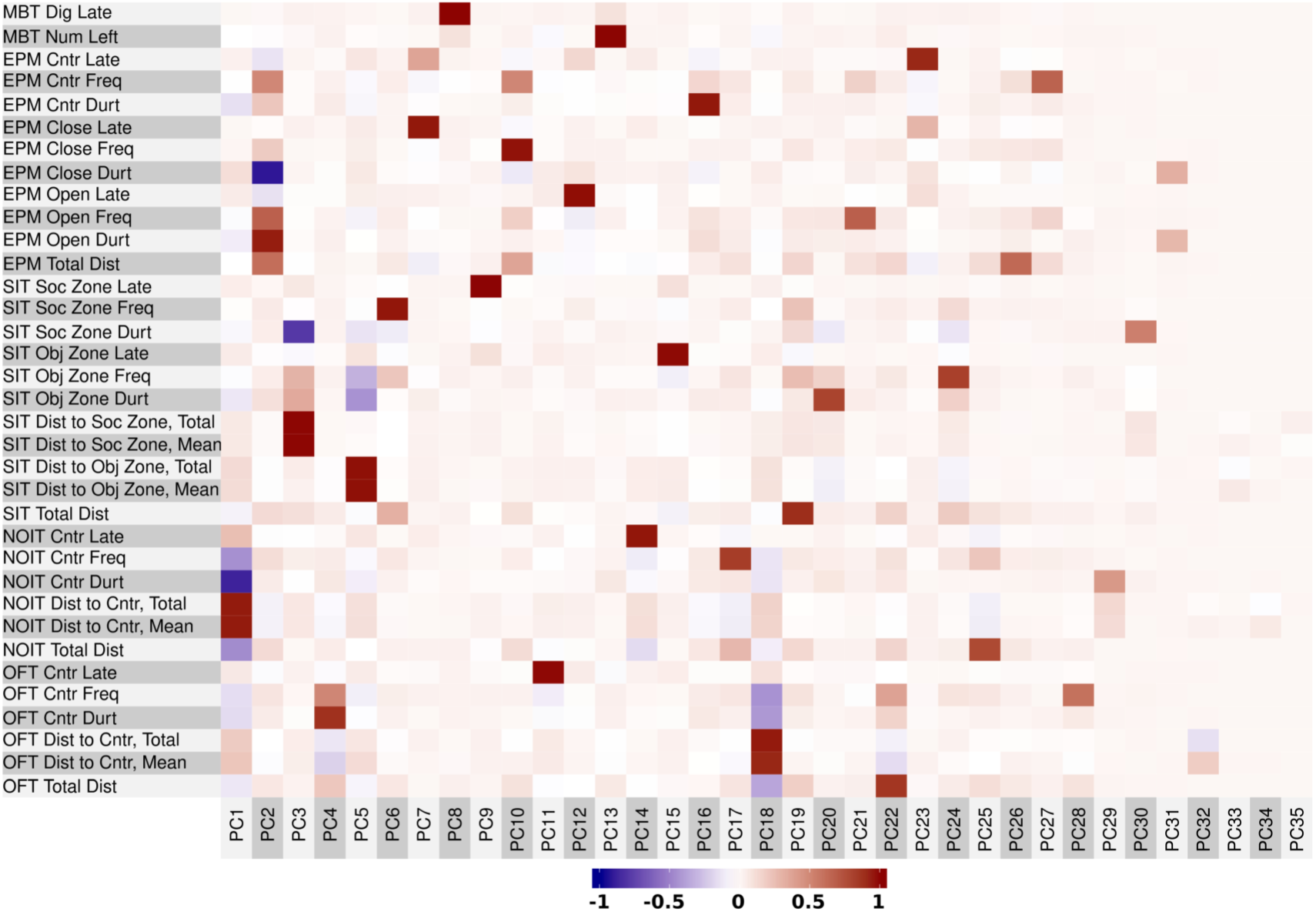
Variable loading for principal component analysis. Principal component (PC) analysis followed by a varimax rotation procedure were conducted on behavioral measures obtained from the five non-nicotine-related behavioral tests. The loading of each PC was plotted as a heatmap.

Table 7 summarizes the results of multiple regression on nicotine infusion during the fixed ratio and progressive ratio sessions and the number of licks on the active spout during context cue-induced reinstatement of drug-seeking. In general, 10–20% of the variance in these measures were explained by approximately 5-6 significant PCs. Notably, PCs 5, 6, 19, & 20 which had strong loadings on the social interaction test, and PCs 18, 21 & 22, which had strong loadings on anxiety-related traits, had significant contributions to several measures of nicotine intake and reinstatement in both sexes. In addition, PCs 2 and 3, which loaded onto anxiety and social behavior, respectively, were significant predictors of nicotine infusion in males. Details for these analysis are provided in Tables S12 – S19. The analysis presented above used PCs generated using both male and female data. We also conducted regression using sex-specific PCs and the amounts of variance explained were very similar (data not shown).

**Table 7.** Summary of principal component regression on nicotine self-administration

| Phenotype | Sex | Significant PCs | R-squared | F | p |
| --- | --- | --- | --- | --- | --- |
| Nicotine infusion first 3 FR sessions | F | **6**, 7, 12, 17, <u>18</u>, <u>22</u> | 0.16 | 4.0e-07 |  |
| Nicotine infusion first 3 FR sessions | M | **5**, <u>18</u>, **19**, <u>21</u>, 23, 25 | 0.14 | 3.9e-05 |  |
| Nicotine infusion last 3 FR sessions | F | 12, **19**, 31 | 0.09 | 0.0004 |  |
| Nicotine infusion last 3 FR sessions | M | 1, 2, 3, **5**, **6**, 13, 15, 17, **20**, 30 | 0.19 | 2.2e-07 |  |
| Nicotine infusion PR session | F | 4, **5**, **6**, <u>18</u>, <u>22</u> | 0.16 | 0.0002 |  |
| Nicotine infusion PR session | M | 1, **5**, **20**, <u>21</u>, <u>22</u>, 31 | 0.19 | 3.6e-06 |  |
| Reinstatement, active lick | F | **6**, **19** | 0.10 | 0.002 |  |
| Reinstatement, active lick | M | **5**, 14, 17, **20**, <u>21</u>, 23 | 0.17 | 2.9e-05 |  |

The social behavior and anxiety associated PCs that frequently significant for nicotine traits were highlighted in bold or underlined, respectively.

### Correlation between nicotine metabolism and self-administration

We measured the level of serum cotinine in 92 rats that received forced nicotine injections one day after the progressive ratio test. Standard curves were fitted using four parameters and *R*^2^s were > 0.99. We found that there was no correlation between cotinine level and the amount of self-administered nicotine during the last 3 nicotine IVSA sessions (r = 0.004, p = 0.97, Figure 5).

**Figure 5.**
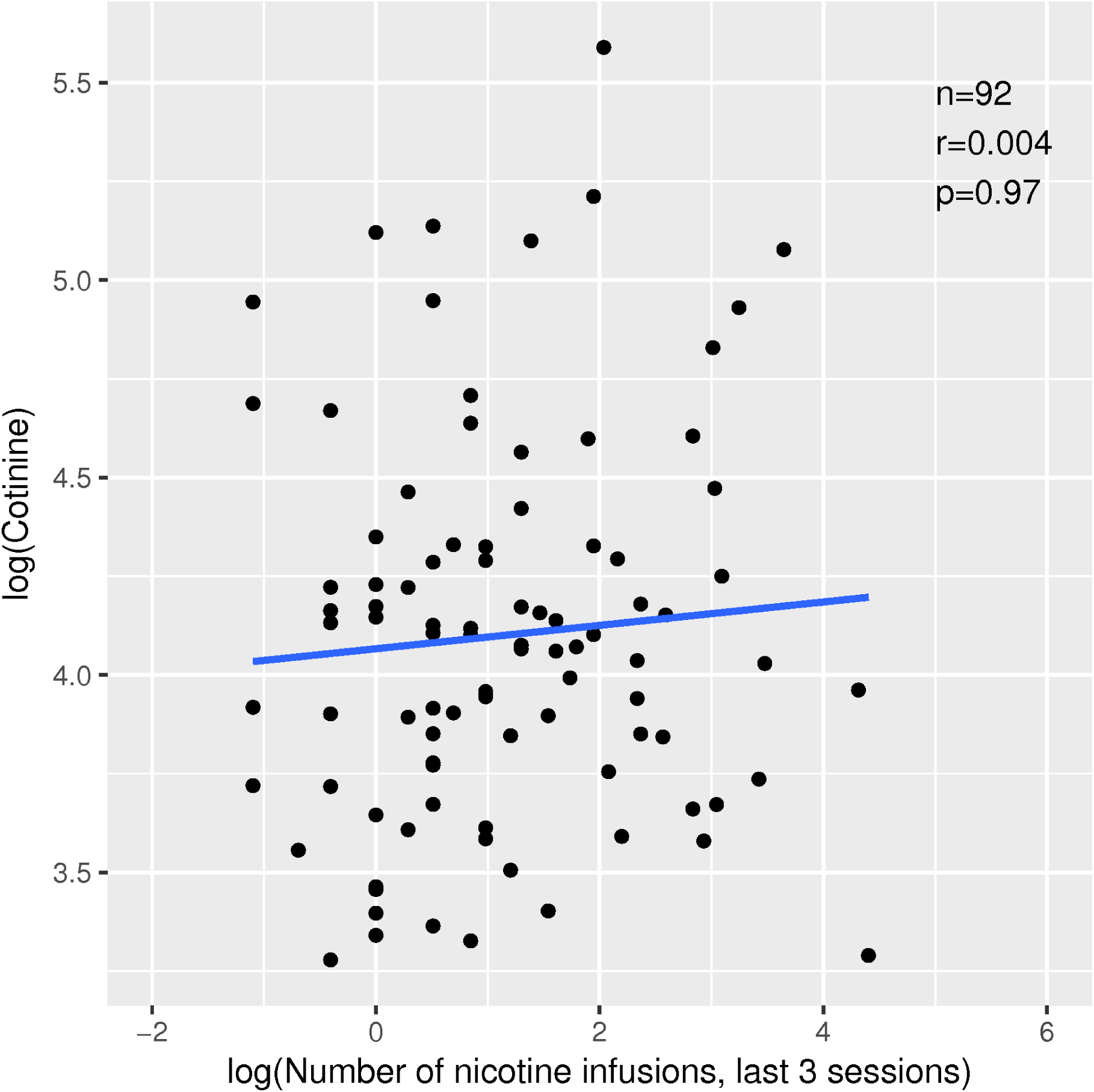
Lack of correlation between nicotine metabolism and nicotine self-administration. Rats received 5 intravenously nicotine injections (30 *μ* g/kg) one day after the progressive ratio test. Blood samples were withdrawn 3 h after the last injection. Plasma cotinine levels were measured using a rat ELISA assay.

## Discussion

This is an interim report of our ongoing genome-wide association study on socially acquired nicotine self-administration. Each of the ~ 800 adolescent heterogeneous stock rats was first tested on five behavioral tests, including OFT, NOIT, SIT, EPM, and MBT. Rats were then implanted with a jugular catheter and trained on the socially acquired nicotine IVSA protocol. DNA samples were collected from each rat and are being analyzed. Our current sample size is approximately half of our goal. However, these data already provide numerous interesting findings, such as 243 significant correlations between behavioral tests (0.07 < |*r*| < 0.40) and 16 significant sex differences in behavioral measures. We also performed a principal component regression analysis and found that ~ 10 – 20% of the variance in nicotine self-administration can be explained by PCs with strong loading of measures of social interaction and anxiety. Lastly, we found that there was no correlation between nicotine intake and nicotine metabolism.

Our study used an outbred population of HS rats. Because of the genetic diversity of this population^28^, we expect there to be large variations in behavioral measures. This indeed was the case (Tables 2–6, Figures 2–3). Because the same rats were tested in all of the behavioral tests, we also expected measures influenced by the same genetic factors to be correlated. As shown in Figure 1, we found 354 significant correlations among measures of non-nicotine phenotypes. As expected, many (111) of these were between measures of the same behavioral test (Tables S1–S5). Most interestingly, we also found 243 significant correlations that were between measures of different behavior tests (Tables S6–S8). The finding of a large number of relatively low coefficient (e.g., *r* < 0.4) but statistically highly significant correlations between behavioral tests is in agreement with other behavioral studies that had similar sample sizes (e.g. Lopez-Aumatell et al.^29^, n = 787). A likely explanation for these correlations was that they were controlled by the same behavioral process and thus were influenced by the same genetic factors.

The NOIT had some of the strongest cross-test correlations. For example, the correlation of the mean distance to the center zone between OFT and NOIT tests was 0.33. The center zone frequencies and durations were also correlated between these two tests (r = 0.27 for both, Table S6). Although this could partically be explained by these two tests shared the same arena, EPM close arm duration, an indicator of anxiety obtained in a different experimental setup, was also correlated with the distance to the center zone measured in the NOIT (r = 0.22, Table S6). These findings suggested that although rats were habituated for 1 h in the arena on the first day of the test, interacting with a novel object in the center of the same arena the next day nevertheless induced a similar approach/avoidance conflict. On the other hand, compared to the OFT, rats entered the center zone with significantly shorter latency, visited the center with higher frequency, spent more time in the center zone during the NOIT, thus indicating that the NOIT indeed measured the natural exploratory behavior^30^ of these HS rats.

Another group of interesting correlations was that between OFT and EPM. Both these tests were commonly used to measure anxiety-like behavior. We found that OFT center frequency was correlated with EPM open arm duration (r = 0.22) and EPM close arm duration (r = -0.15). Distance to the center zone in the OFT was correlated with EMP close (r = 0.16) and open (r = -0.23) arm duration. Also, OFT center latency was correlated with EPM open arm latency (r = 0.08), and open arm frequency was correlated with OFT center frequency (r = 0.09). Because of the relatively small correlation coefficients, these data thus supported the notion that EPM and OFT were measuring different aspects of the anxiety phenotype^31^.

The third group of intriguing correlations involved total travel distance. Correlations of total travel distance between the four tests where they were measured were among the strongest in this dataset (0.29 < *r* < 0.39). Because freezing is a common response of rodents to fear and anxiety^32^, one likely explanation for this correlation is that these tests all invoked generalized anxiety.

This postulate was further supported by the strong negative correlation between total distance and close arm duration in the EPM test (r = -0.62).

There were also tests that had little correlation with others, such as those measuring the social zone in SIT and the MBT. These suggested that social interaction and marble bury measured aspects of rat behavioral repertoire that were not related to anxiety or fear.

We found many sex differences in behavior tests. Overall, adolescent male HS rats have heightened levels of anxiety than do females. Compared to females, the males stayed further away from the center of the arena in the OFT and NOITs, spent less time in the center zones, and made fewer visits to the center zone (Table 2 and Table 3); they also entered the open arms after longer delays, paid fewer visits to the open arms, and spent less time exploring the open arms in the EPM test (Table 5). These sex differences in the HS population were in agreement with those reported by Lopez-Aumatell et al.^29^ and by us^33^ for the adult HS rats.

We also found a strong sex difference in nicotine IVSA. Without exceptions, females had more robust nicotine intake behavior than males: they obtained more nicotine during the first and the last three fixed-ratio sessions (Figure 3 A and B), they had higher breakpoints in the progressive ratio session and showed a greater number of licks on the active spout in the reinstatement test (Figure 3D). These differences are in agreement with our previous finding in HS adults^33^. This overall sex difference potentially indicated enhanced vulnerability of females to the reinforcing properties of nicotine in a social environment. Alternatively, it could suggest that the heightened anxiety in males could potentially have a protective effect against nicotine intake. This relationship could be specific to the HS rats because prior studies have shown that the relationship between nicotine consumption and anxiety depends on both the age and the genetic background of rodents^34^.

When compared to our previous finding using adult HS rats^33^, the data presented here also indicated that there was a large difference in these behaviors when measured at different ages. For example, the center duration for the adults in the OFT was almost three times that of the adolescents for both males and females. Similarly, the percent time spent by adult HS rats in exploring the open arm in the EPM test was twice that of the adolescents for both males and females. Although these data were collected several years apart, they were collected using genetically similar rats, the same equipment and analyzed using the same software. Given the large magnitude of the difference, it is very likely that adolescent HS rats have higher levels of anxiety than adults.

The main phenotype of this study is socially acquired nicotine IVSA. We previously reported a large sex difference in nicotine intake in 100 adult HS rats using a protocol where a demonstrator rat consuming the flavor cue was present during the IVSA sessions^33^. The interpretation of these results were complicated by the finding of a large sex difference in the behavior of the demonstrator rats, where the females consumed more flavor cue but had fewer social interactions (i.e., nose poke into the divider) than the males. Because of the social nature of this IVSA method, the effects of the demonstrator rats’ behavior and the genetic factors of the HS rats on nicotine intake were confounded. More recently, we have reported that carbon disulfide, a chemical component of mammalian breath that mediates social transmission of food preference^35^ was also effective in supporting nicotine IVSA with a flavor cue^18^. With this new approach, the effect of the demonstrator rat was recapitulated by adding a component of their breath into the flavor cue. This effectively removed the behavior of the demonstrator rat as a confounding factor while retained the social nature of our nicotine IVSA method. This social nature was further validated by regression analysis showing that factors corresponding to measures of social behaviors were among the strongest predictors of nicotine intake and reinstatement of nicotine-seeking in this model (see below).

In general, the average nicotine intake in the adolescent HS rats was lower than those we previously reported for the Sprague-Dawley rats^17^, which is also an outbred population. This difference is likely due to several founding strains of the HS population, including Fisher 344 and Wistar-Kyoto, which show low nicotine intake when tested using the socially acquired protocol^20^ or the lever pressing protocol^36,37^. To better appreciate the variation in self-administration data, we split the data using the median nicotine intake for the 10 fixed-ratio sessions. In both females and males, rats with higher nicotine intake showed a significantly greater number of licks on the active spout compared to the inactive spout, while the opposite pattern was seen for those with lower nicotine intake (Figure 2). Figure 3C and 3D show large variations in the breakpoint in progressive ratio test and the reinstatement test. These data indicated that the genetic diversity of this population is a significant contributor to the divergent nicotine IVSA behavior and further support the use of this population for mapping genetic factors contributing to nicotine-taking behavior. Also, the reinstatement data showed that both female and male rats licked significantly more on the active spout than on the inactive spout and that there was a significant sex difference in the number of licks on the active spout but not on the inactive spout (Figure 3 D, data include both the low and high nicotine intake groups). These results confirmed that despite the relative low nicotine intake, these rats did develop a preference for nicotine.

Many genetic studies on drug addiction traits have found protection genes rather than vulnerability genes. For example, the CHRNA5 gene associated with smoking cessation and lung cancer is responsible for the aversive effect of nicotine^38^. Similarly, the alcohol dehydrogenase and aldehyde dehydrogenase variants associated with drinking provide protective effects^39^. In animal studies, we sequenced the nucleus accumbens shell transcriptome of 10 isogenic rat strains and found 12 gene clusters associated with nicotine intake^36^. Nine of these clusters had negative correlation coefficients, suggesting they play protective roles^40^. Similarly, because of the relatively low nicotine intake at the population level (with many individuals showing enhanced nicotine intake), we anticipate many of the future genetic findings from these rats are likely going to be genes that provide a protective effect on smoking in the social environment.

The most intriguing insights of these studies came from the principal component regression analysis. As shown in Table 7 and Tables S12 – S19, several PCs with strong loadings on the SIT were significant predictors for several nicotine phenotypes. For example, PC 5, with loadings on the distance to the object zone in the SIT, was significant for nicotine infusion during the first and last three fixed ratio sessions, the progressive ratio session, and reinstatement in males, as well as for nicotine infusion in the progressive ratio session in females. Similarly, PC 6, with strong loading on social zone frequency, was significant for nicotine infusion in the first three fixed-ratio sessions, the progressive ratio session, and reinstatement in females and the last three fixed-ratio sessions in males. Further, PC 19 and 20, with strong loadings on total travel distance and object zone duration in SIT, respectively, were each significant for three nicotine phenotypes. These results were similar to our previous finding in adult HS rats that PCs representing social phenotypes are the strongest predictors for nicotine phenotypes in the socially acquired nicotine IVSA model, despite the fact that the nicotine IVSA presented here were conducted using the modified method where carbon disulfide was used to replace live demonstrator rats. These regression results and their agreement with previous findings further validated the carbon disulfide model. In addition, anxiety-related traits are significant predictors for nicotine phenotypes. For example, PC 18 and 22 with strong loading on several OF measures, and PC 21 with primary loading on open arm frequency in the EPM test, were each significant for three nicotine-related measures.

Regression analysis also indicated strong sex differences. In general, more unique PCs were significant for the males (18) than for the females (10) (Table 7). For example, the first three PCs, with loading on the NOIT, EPM, and SIT, respectively, were all significant for the males but not for the females. Further, except nicotine intake during the first three sessions, more variance was explained in the regression analysis for males than for females. Considering that females consumed more nicotine than males (Figure 2), we speculate that the reinforcing effect of nicotine, which is not included in the regression analysis, is potentially stronger in females than in males, while anxiety and social learning, which was represented by the PCs in the regression analysis, play more prominent roles in the nicotine intake behavior of adolescent males.

One potential contributor to individual variations in drug-taking behavior is drug metabolism. The limited strain-specific data suggested that metabolism is not likely a major factor in rat nicotine IVSA. For example, nicotine metabolism is similar between Fisher 344 and Sprague-Dawley rats^41^. While Sprague-Dawley rats readily acquire nicotine IVSA, Fisher 344 rats do not^36,36^. We measured the level of cotinine, the major metabolite of nicotine, in the first 100 rats of this study after they completed nicotine IVSA. We found no evidence that cotinine level was correlated with nicotine intake (Figure 5). Therefore, it is unlikely nicotine metabolism is a significant contributor to nicotine intake in our model.

In summary, our data showed large variation in anxiety, social, and nicotine-driven behavior in adolescent HS rats. A large number of correlations between these behaviors suggested that they are likely influenced by the same genetic factors. Further, our regression analysis indicated that anxiety and social phenotypes contribute to approximately 10-20% of the variances in both males and females. We also found many sex differences in this analysis, indicating a high probability of discovering sex-specific genetic mechanisms for nicotine intake in future studies.

## Methods

### Animals

A total of 8 batches of Heterogeneous Stock rats (NMcwi:HS) rats were processed between 2014-2017. Each batch started with 25 male and 25 female breeders transferred from the Medical College of Wisconsin to the University of Tennessee Health Science Center. After a two week quarantine period, these rats were transferred to a reversed 12:12 h light–dark cycle (lights off at 9:00 AM) housing room. Breeding pairs were assigned to maximize genetic diversity of the population. Litters were culled to a maximal of 8 pups to ensure standard nutrition. Rats were weaned on postnatal day (PND) 21. A radio frequency identification chip (RFID) was inserted subcutaneously when rats were weaned. Two male and two female rats per litter were used for behavioral studies. Sprague-Dawley rats (20 for each sex, purchased from Harlan Laboratories, Madison, WI) were used as the stimulus rats in the social interaction test. An additional 50 female and 50 male Sprague-Dawley rats were used as demonstrators used for context induced reinstatement test. Standard rat chow and water were provided *ad libitum*. All rats were group housed with 2-4 same sex peers throughout the experiments to avoid social isolation. All procedures were conducted in accordance with the NIH Guidelines concerning the Care and Use of Laboratory Animals, as approved by the Institutional Animal Care and Use Committee of the University of Tennessee Health Science Center.

### Study Design

All HS rats born in UTHSC were adolescents (PND 32) when tests began. Each HS rat was first tested in a series of behavioral tests, one test per day, conducted in the dark-phase of the light cycle (9AM - 4PM). These tests were conducted in the following sequence: OFT, NOIT, SIT, EPM and MBT.

The first three tests were conducted in the same open field and recorded using the same video capture system. The sequence and procedures of these tests were adapted from two articles^42,43^. The schedule for behavioral tests are provided in Table 1

### Open field test

The open field test has been used to study anxiety in animal models. Two OFT chambers constructed using black acrylic glass, measuring 100 cm (L) x 100 cm (W) x 50 cm (H), were placed side by side. The floors were wood boards painted with black or white acrylic paint (ART-Alternatives, ASTM D-4236, Emeryville, CA, USA) to contrast the coat of the animals. The test chambers were illuminated by a long-range, 850-nm infrared illuminator (LIR850-70, LDP LLC, Carlstadt, NJ) located 160 cm above the center of the two test chambers. No source of visible light was present during behavioral testing with the exception of a flat panel monitor (Dell 1908FP). A digital camera (Panasonic WV-BP334) fitted with an 830 nm infrared filter (X-Nite830-M37, LTP LLC, Carlstadt, NJ) and located next to the infrared light source was used to record the behavior of the rats. All rats were released at the same corner of the test chamber and data were collected for 1 h on day 1 and 20 min on day 2 (novel object) and day 3 (social interaction). Ethovision^®^ XT video tracking system (Version 4.0, Noldus Information Technology, The Netherlands) was used to obtain the total distance traveled and the duration that the animal was present in the center of the test field (a circular region with a diameter of 20 cm).

### Novel object interaction test

Novelty seking has often been found to be associated with drug abuse in both animal model and human subjects^44^. In this test, the novel object was a cylindrical cage constructed using 24 aluminum rods (30 cm in length) that were spaced 1.7 cm apart. The bottom and top of the cage (15 cm in diameter) were manufactured using Makerbot Replicator 2 (Makerbot Industries, Brooklyn, NY). The design can be downloaded from https://www.github.com/chen42/vighrdesigns. On novel object was placed into the center of each open field before testing. Other aspects of the test were the same as open field test with the exception that behavior was only recorded for 20 min.

### Social interaction test

This modified social interaction test compares the preference of a test subject for a sex and weight matched rat against a cylindical cage. The test arena were measured as 100 cm (L) × 60 cm (W) × 50 cm (H). Two cylindrical cages described above were placed ~ 30 cm away from the walls on opposite sides (i.e., similar to the arrangement commonly used in the three chamber test). A randomly selected stimulus Sprague-Dawley rat of the same sex as the HS test rat was placed into one assigned cage (kept the same throughout the experiment) 5 mins before the HS subject rat was placed into the open field. The stimulus and test rats were never housed together and thus were unfamiliar to each other. No social isolation was conducted on either rat. Each stimulus rat was used once per day. The test duration was 20 min.

### Elevated plus maze test

The elevated plus-maze was a test for measuring anxiety^45^ in rodents. The maze was constructed using black acrylic glass. The floors of the maze were covered by wood boards painted with black or white acrylic paint. The platform was 60 cm above the floor, with all four arms measuring 12 cm (W) × 50 cm (L). The two opposing closed arms had walls measuring 30 cm (H). Rats were placed into the center of the maze facing the closed arm. The behavior of the rat was recorded for 6 min using the digital video system described above. Ethovision was used to extract the duration, frequency and frequency each rat spent in each arm.

### Marble bury test

The Marble bury test was often used to model obsessive-compulsive behavior^46^. It was conducted using a regular rat housing cage, measuring 47 cm (L) × 24 cm (W) × 15 cm (H), filled with 5 cm (depth) of corn cob bedding. Twenty round dark-blue colored glass marbles (diameter 2.54 cm) were distributed evenly in a 5 × 4 grid. The cage and bedding were used for a maximum of 6 same sex rats. Rats were patted for ten times on their heads before they were placed in the center of the cage. A filter top cover was then placed on the cage. The latency to dig the bedding (seconds) was recorded using a stop watch, with a maximum allowed time of 3 minutes. Room lights were then turned off, and the technician left the room. After a total of 20 minutes, rats were removed from the test cage and a photo of the cage was taken. The number of marbles left on top of the bedding were counted manually from the photograph. Only marbles with more than half of its diameter visible were counted.

### Nicotine self-administration in social context

Nicotine IVSA was conducted according to our published protocol^18^ with some modifications. Rats were implanted with jugular catheters constructed using Micro-Renathane tubing (MRE037, Braintree Scientific Inc., Braintree, MA) under isoflurane anesthesia. The tubing exited from the back of the rat through a 3D printed implant^47^. Carprofen (5 mg/kg, s.c.) was given immediately before surgery for analgesia. After three days of recovery from the surgery, rats were given access to nicotine IVSA for 2.5 h per day for 10 days in the dark-phase of the light cycle. Experiments were conducted 5 days per week. Training was continued during the weekends, if less than 3 sessions had been completed by a rat. Each operant chamber (Med Associates, St Albans, VT) contained a divider with six equally spaced circular holes that allow orofacial contacts between rats. The operant chambers were located in sound attenuating chambers and each contained two drinking spouts fitted on the same wall. Two syringe pumps were placed outside of each sound attenuating chamber; one delivered i.v. nicotine through a swivel located on top of the chamber and the other delivered the flavor cue to the active spout. Each spout was connected to a contact lickometer controller that records the number and the timing of licks. IVSA was conducted using a fixed-ratio 10 schedule with 20 s timeout period (FR10TO20). Thus, 10 licks on the active spout activated the simultaneous delivery of a 60 *μ* l flavor cue, and an i.v. infusion (nicotine free base, 15 *μ* g/kg or saline). The flavor cue contained saccharin (0.2%), unsweetened grape Kool-Aid (0.1%), and carbon disulfide (500 ppm), which we have shown to support socially acquired nicotine self-administration in the absence of a demonstrator rat^18^. Licks on the inactive spout had no programmed consequence. Licks during the timeout period had no consequences but were recorded. No audio or visual cue was used. Rats were not food or water deprived. Nor did rats receive operant training or priming nicotine injections before the initiation of the SA sessions.

One progressive ratio test was conducted on the 11th session. The number of licks to obtain a subsequent infusion was determined using the exponential formula 5*e*^0.2×*injections*^ – 5, such that the required responses per injection were as follows: 1, 2, 4, 9, 12, 15, 20, 25, 32, 40, 50, etc.^48^ The sessions ended after 20 min of inactivity. The final ratio completed is the breakpoint. The patency of the jugular catheters was tested using a fast-acting anesthetic, methohexital (0.1 ml, 10 mg/ml), at the end of the progressive ratio session for each rat. Rats without functional catheters were excluded from the analysis.

### Context induced reinstatement of drug-seeking

Context extinction and reinstatement procedures were conducted after the progressive ratio sessions. Context extinction chambers were different from the IVSA operant chamber in many aspects, including the absence of the divider, flat floor, and distinct audiovisual cues. There were two clean dry spouts in the chamber and the number of licks were recorded but had no programmed consequence. Licking data were collected using a capacitive touch sensor connected to a single board computer (Raspberry PI). A detailed description of the setup and software are available in our GitHub repository (https://github.com/chen42/openbehavior) (under Extinction). The self-administering rats were placed in the extinction environment for 1 h/d until the lick on the “active” spout was reduced to less than 50 for two consecutive sessions.

Reinstatement of drug-seeking behavior was tested the day after the last context extinction session. The reinstatement rat and a demonstrator rat of the same sex were placed in the nicotine IVSA chamber on the opposite side of the divider. The demonstrator rat had free access to the flavor cue, while the reinstatement rat did not. The number of licks on the spouts and nose pokes into the divider from both rats were recorded for 1 h. Licking from the reinstatement rat had no programmed consequences.

### Blood cotinine level

We measured cotinine, the major nicotine metabolite, one day after nicotine IVSA was finished in the first 100 rats. Rats were placed in the operant chambers and received 5 injections of nicotine (30 *μ* g/kg) through jugular catheter at 30 s intervals. Blood samples were withdrawn from the jugular catheter 3 h after the last injection. This time point was chosen because our past data indicated 3 h was the peak of blood cotinine concentration using this protocol. Plasma cotinine levels were measured using a rat ELISA assay (Calbiotech).

### Statistical analysis

Data were analyzed using the R language and environment for statistical computing and graphics. Data were presented as mean ± SD. Number of licks and infusions were analyzed using repeated-measures ANOVA with spout and session treated as within-subject variables. Paired t-tests were used to compare the number of licks on the two spouts during reinstatement. T-tests were used to compare the sex differences on the means. Effect size (Cohen’s d) was calculated using pooled SD. Correlation between behavioral test scores were calculated using Pearson’s r (psych package). Multiple testing was controlled using the Holm’s method. Because of the multicollinearity in the dataset, we used the principal component analysis (PCA) to obtain orthogonal representation of the data using the “principal” function available from the psych package. These PCs were then tested as predictors for nicotine intake in a multiple regression analysis using the “lm” function. The “step” function was used to select variables based on the Akaike information criterion. Statistical significance was assigned when p < 0.05.

### Data Availability

The datasets generated during and/or analysed during the current study are available from the corresponding author on reasonable request.

## Acknowledgements

The authors thank Pawandeep Kaur, Yanyan Lin, Shen Jie, and Hong Xia for their technical assistance in conducting these experiments. Funding was provided by NIDA Grant DA-026894 (AP, HC, LW).

## Author contributions statement

H.C., A.P. and L.W conceived of the experiments, T.W. and W.H. conducted the experiments and drafted the manuscript, T.W., A.C., O.P. and H.C. analyzed the results. All authors reviewed the manuscript.

## References

1. Eissenberg, T. & Balster, R. Initial tobacco use episodes in children and adolescents: current knowledge, future directions. Drug Alcohol Depend 41–60 (2000).

2. Greenlund, K., Johnson, C., Webber, L. & Berenson, G. Cigarette smoking attitudes and first use among third-through sixth-grade students: the bogalusa heart study. Am J Public Heal. 87, 1345–1348 (1997).

3. White, V., Byrnes, G., Webster, B. & Hopper, J. Does smoking among friends explain apparent genetic effects on current smoking in adolescence and young adulthood? Br J Cancer 98, 1475–1481 (2008).

4. Memetovic, J., Ratner, P., Gotay, C. & Richardson, C. Examining the relationship between personality and affect-related attributes and adolescents’ intentions to try smoking using the substance use risk profile scale. Addict Behav 56, 36–40 (2016).

5. Banzer, R. et al. Factors associated with different smoking status in european adolescents: results of the seyle study. Eur Child Adolesc Psychiatry 26, 1319–1329 (2017).

6. Barrera, D. Gender differences in the transmission of smoking from filipino parents to their offspring: The role of parenting, school climate, and negative emotions. Subst Use Misuse 52, 1439–1448 (2017).

7. Moxley, R. & Olson, L. Clinical evaluation of transmissible gastroenteritis virus vaccines and vaccination procedures for inducing lactogenic immunity in sows. Am J Vet Res 50, 111–118 (1989).

8. Galera, C. et al. Stress, attention deficit hyperactivity disorder (adhd) symptoms and tobacco smoking: The i-share study. Eur Psychiatry 45, 221–226 (2017).

9. Pergadia, M., Heath, A., Martin, N. & Madden, P. Genetic analyses of dsm-iv nicotine withdrawal in adult twins. Psychol Med 36, 963–972 (2006).

10. Carmelli, D., Swan, G., Robinette, D. & Fabsitz, R. Genetic influence on smoking–a study of male twins. N Engl J Med 327, 829–833 (1992).

11. Hall, W., Madden, P. & Lynskey, M. The genetics of tobacco use: methods, findings and policy implications. Tob Control. 11, 119–124(2002).

12. Li, M., Cheng, R., Ma, J. & Swan, G. A meta-analysis of estimated genetic and environmental effects on smoking behavior in male and female adult twins. Addict. 98, 23–31 (2003).

13. Tobacco and Genetics Consortium. Genome-wide meta-analyses identify multiple loci associated with smoking behavior. Nat Genet. 42, 441–447 (2010).

14. Saccone, N. et al. The chrna5-chrna3-chrnb4 nicotinic receptor subunit gene cluster affects risk for nicotine dependence in african-americans and in european-americans. Cancer Res 69, 6848–6856 (2009).

15. Thorgeirsson, T. et al. A variant associated with nicotine dependence, lung cancer and peripheral arterial disease. Nat. 452, 638–642 (2008).

16. Thorgeirsson, T. et al. Sequence variants at chrnb3-chrna6 and cyp2a6 affect smoking behavior. Nat Genet. 42, 448–453 (2010).

17. Chen, H., Sharp, B., Matta, S. & Wu, Q. Social interaction promotes nicotine self-administration with olfactogustatory cues in adolescent rats. Neuropsychopharmacol. 36, 2629–2638 (2011).

18. Wang, T. & Chen, H. Carbon disulfide mediates socially-acquired nicotine self-administration. PLoS One 9, e115222 (2014).

19. Wang, T., Han, W. & Chen, H. Socially acquired nicotine self-administration with an aversive flavor cue in adolescent female rats. Psychopharmacol. (Berl) 233, 1837–1844(2016).

20. Han, W., Wang, T. & Chen, H. Social learning promotes nicotine self-administration by facilitating the extinction of conditioned aversion in isogenic strains of rats. Sci Rep 7, 8052 (2017).

21. Hansen, C. & Spuhler, K. Development of the national institutes of health genetically heterogeneous rat stock. Alcohol Clin Exp Res 8, 477–479.

22. Mott, R., Talbot, C., Turri, M., Collins, A. & Flint, J. A method for fine mapping quantitative trait loci in outbred animal stocks. Proc Natl Acad Sci U S A 97, 12649–12654 (2000).

23. Baud, A. et al. Combined sequence-based and genetic mapping analysis of complex traits in outbred rats. Nat Genet. 45, 767–775 (2013).

24. Solberg Woods, L., Holl, K., Tschannen, M. & Valdar, W. Fine-mapping a locus for glucose tolerance using heterogeneous stock rats. Physiol Genomics 41, 102–108 (2010).

25. Solberg Woods, L. et al. Fine-mapping diabetes-related traits, including insulin resistance, in heterogeneous stock rats. Physiol Genomics 44, 1013–1026 (2012).

26. Holl, K. et al. Heterogeneous stock rats: a model to study the genetics of despair-like behavior in adolescence. Genes Brain Behav (2017).

27. Keele, G. et al. Genetic fine-mapping and identification of candidate genes and variants for adiposity traits in outbred rats. Obes. (Silver Spring) 26, 213–222 (2018).

28. Woods, L. & Mott, R. Heterogeneous stock populations for analysis of complex traits. Methods Mol Biol 1488, 31–44 (2017).

29. Lopez-Aumatell, R. et al. Fearfulness in a large n/nih genetically heterogeneous rat stock: differential profiles of timidity and defensive flight in males and females. Behav Brain Res 188, 41–55 (2008).

30. Blaser, R. & Heyser, C. Spontaneous object recognition: a promising approach to the comparative study of memory. Front Behav Neurosci 9, 183 (2015).

31. Ramos, A., Pereira, E., Martins, G., Wehrmeister, T. & Izídio, G. Integrating the open field, elevated plus maze and light/dark box to assess different types of emotional behaviors in one single trial. Behav Brain Res 193, 277–288 (2008).

32. Canteras, N., Resstel, L., Bertoglio, L., Carobrez, A. P. & Guimarães, F. Neuroanatomy of anxiety. Curr Top Behav Neurosci 2, 77–96 (2010).

33. Wang, T. et al. Propensity for social interaction predicts nicotine-reinforced behaviors in outbred rats. Genes Brain Behav 13, 202–212 (2014).

34. Wilking, J. et al. Comparison of nicotine oral consumption and baseline anxiety measures in adolescent and adult c57bl/6j and c3h/ibg mice. Behav Brain Res 233, 280–287 (2012).

35. Galef, B., Mason, J., Preti, G. & Bean, N. Carbon disulfide: a semiochemical mediating socially-induced diet choice in rats. Physiol Behav 42, 119–124 (1988).

36. Chen, H., Hiler, K., Tolley, E., Matta, S. & Sharp, B. Genetic factors control nicotine self-administration in isogenic adolescent rat strains. PLoS One 7, e44234 (2012).

37. Sharp, B. et al. Gene expression in accumbens gaba neurons from inbred rats with different drug-taking behavior. Genes Brain Behav 10, 778–788 (2011).

38. Fowler, C., Lu, Q., Johnson, P., Marks, M. & Kenny, P. Habenular alpha5 nicotinic receptor subunit signalling controls nicotine intake. Nat. 471, 597–601 (2011).

39. Edenberg, H. The genetics of alcohol metabolism: role of alcohol dehydrogenase and aldehyde dehydrogenase variants. Alcohol Res Heal. 30, 5–13 (2007).

40. Chen, H., Luo, R., Gong, S., Matta, S. & Sharp, B. Protection genes in nucleus accumbens shell affect vulnerability to nicotine self-administration across isogenic strains of adolescent rat. PLoS One 9, e86214 (2014).

41. Kyerematen, G., Owens, G., Chattopadhyay, B., deBethizy, J. & Vesell, E. Sexual dimorphism of nicotine metabolism and distribution in the rat. studies in vivo and in vitro. Drug Metab Dispos 16, 823–828.

42. Green, M., Barnes, B. & McCormick, C. Social instability stress in adolescence increases anxiety and reduces social interactions in adulthood in male long-evans rats. Dev Psychobiol 55, 849–859 (2013).

43. Vetter-O’Hagen, C. & Spear, L. The effects of gonadectomy on sex-and age-typical responses to novelty and ethanol-induced social inhibition in adult male and female sprague-dawley rats. Behav Brain Res 227, 224–232 (2012).

44. Wingo, T., Nesil, T., Choi, J. & Li, M. Novelty seeking and drug addiction in humans and animals: From behavior to molecules. J Neuroimmune Pharmacol 11, 456–470 (2016).

45. Lezak, K., Missig, G. & Carlezon, W. Behavioral methods to study anxiety in rodents. Dialogues Clin Neurosci 19, 181–191 (2017).

46. Albelda, N. & Joel, D. Animal models of obsessive-compulsive disorder: exploring pharmacology and neural substrates. Neurosci Biobehav. Rev 36, 47–63 (2012).

47. Wang, T., Wang, B. & Chen, H. Menthol facilitates the intravenous self-administration of nicotine in rats. Front Behav Neurosci 8, 437 (2014).

48. Richardson, N. & Roberts, D. Progressive ratio schedules in drug self-administration studies in rats: a method to evaluate reinforcing efficacy. J Neurosci Methods 66, 1–11 (1996).

